# Mechanosensitive regulation of FGFR1 through the MRTF-SRF pathway

**DOI:** 10.1101/782243

**Authors:** Jip Zonderland, Silvia Rezzola, Lorenzo Moroni

## Abstract

Controlling basic fibroblast growth factor (bFGF) signaling is important for both tissue-engineering purposes, controlling proliferation and differentiation potential, and for cancer biology, influencing tumor progression and metastasis. Here, we observed that human mesenchymal stromal cells (hMSCs) no longer responded to soluble or covalently bound bFGF when cultured on microfibrillar substrates, while fibroblasts did. This correlated with a downregulation of FGF receptor 1 (FGFR1) expression of hMSCs on microfibrillar substrates, compared to hMSCs on conventional tissue culture plastic (TCP). hMSCs also expressed less SRF on ESP scaffolds, compared to TCP, while fibroblasts maintained high FGFR1 and SRF expression. Inhibition of actin-myosin tension or the MRTF/SRF pathway decreased FGFR1 expression in hMSCs, fibroblasts and MG63 osteosarcoma cells. This downregulation was functional, as hMSCs became irresponsive to bFGF in the presence of MRTF/SRF inhibitor. Together, our data show that hMSCs, but not fibroblasts, are irresponsive to bFGF when cultured on microfibrillar susbtrates by downregulation of FGFR1 through the MRTF/SRF pathway. This is the first time FGFR1 expression has been shown to be mechanosensitive and adds to the sparse literature on FGFR1 regulation. These results could open up new targets for cancer treatments and could aid designing tissue engineering constructs that better control cell proliferation.

## Introduction

Guiding cell behavior is a critical aspect for cell-based tissue engineering constructs. Among the different tissue engineering scaffolds, electrospun (ESP) scaffolds are being widely studied and are particularly interesting for defects of limited depth, but with a large surface area. Examples include cartilage repair^[1]^, skin patches^[2]^, corneal regeneration^[3]^, nerve guides^[4]^ and vascular grafts^[5]^, among others.

Electrospinning is a technique where synthetic or natural polymers are dissolved in volatile solvents and deposited onto a collector using a highly charged electric field. While traveling from the needle to the collector, the polymer solution becomes highly unstable and is elongated significantly. The solvents evaporate and the polymer is deposited on the collector, resulting in a highly porous fibrous mesh with individual fiber diameters typically ranging 100nm-10μm, depending on processing parameters.

Sufficient number of cells and the right cell density is highly important for tissue engineering applications, so cell proliferation is a key aspect to control. Several growth factors are well known for their proliferation inducing abilities. Arguably, the most well-studied of these is basic fibroblast growth factor (bFGF). bFGF is known to increase proliferation rates in a wide variety of cell types and has anti-apoptotic effects, while maintaining or enhancing differentiation- and regeneration potential^[6]^.

In solution, bFGF, like most growth factors, is highly unstable and loses activity after 24-48 hours^[6, 7]^. Covalently coupling bFGF to scaffolds has been shown to enhance stability while maintaining signaling activity^[8–10]^. Nur et al. showed that covalently coupled bFGF to electrospun fibers maintained activity for 6 months when stored dry^[10]^. When bFGF was covalently coupled to a heparin-mimicking polymer, it maintained increased stability under normal storage conditions^[9]^, and under several stress conditions such as heat or acidic conditions, as opposed to bFGF in solution^[8]^.

bFGF can bind to 7 FGF receptors (coming from 4 FGFR genes, FGFR1-4); tyrosine kinase receptors that can activate a variety of pathways, including the RAS-MAPK, PI3K-Akt, PLCγ and STAT pathways^[11]^. Using next generation sequencing to analyze 4853 tumors, Helsten T. et al. found aberrations in FGFR’s in 7,1% of all tumors^[12]^. In addition, increased expression of FGFR’s has been correlated with a bad prognosis, increased metastasis and tumor progression in a large variety of cancers^[13–17]^. Indeed, animal studies and clinical trials are currently ongoing to test the effects of FGFR inhibitors on cancer treatment, showing promising initial results^[18–23]^. This highlights the importance of understanding how FGFR expression is regulated. Very little is known about the regulation of any FGFR, while a better understand could advance the understanding of tumor development and open up new therapeutic targets.

Besides the role in cancer development, FGFR’s are also interesting for tissue engineering purposes. FGFR1 and 2 have been shown to be involved in adipo- and osteogeneic differentiation in hMSCs^[24, 25]^. FGFR3 is highly expressed in chondrocytes and is involved in chondrogenesis^[26]^. Only FGFR1 has been shown to be involved in hMSCs proliferation^[27]^, while the other receptors remain unstudied in this regard. For this reason, here we focused on the regulation of FGFR1 expression.

Cells adhere to their surrounding matrix or culture substrate through integrins^[28]^. When enough force can be applied, integrin clusters can bind to the actin cytoskeleton through large protein complexes called focal adhesions^[29]^. On the other end, actin filaments can be attached to other focal adhesions, or to the nucleus^[30]^. Between these attachment points, force can be generated by actin-myosin filaments to generate cellular tension^[31]^. A large variety of cellular processes are regulated by cellular tension, including proliferation^[32–35]^, differentiation^[36–38]^ and migration^[39]^. Different transcription factors have been shown to orchestrate these changes in behaviors, of which serum response factor (SRF) is a well-studied example. When globular actin concentrations are low in the cytoplasm, myocardin related transcription factor (MRTF) A or B enters the nucleus and binds to SRF to start transcribing target genes^[40]^.

Here, we have found that hMSCs don’t respond to soluble or covalently bound bFGF when cultured on microfibrillar substrates such as those created by ESP, while fibroblasts do. hMSCs, but not fibroblasts, downregulate FGFR1 expression when cultured on ESP scaffolds. We show that FGFR1 expression is mechanosensitive and works through actin-myosin tension and the MRTF/SRF pathway. Inhibition of the MRTF/SRF pathway made hMSCs irresponsive to bFGF on tissue culture plastic (TCP) and downregulated FGFR1 in hMSCs, fibroblasts and MG63 osteosarcoma cells.

## Results

### Fibroblasts, but not hMSCs, respond to bFGF functionalized ESP scaffolds

Cell-laden ESP scaffolds have been shown to aid tissue repair by hMSCs^[41–43]^ and fibroblasts^[44–46]^ in vivo. bFGF is known to enhance proliferation while maintaining differentiation potential in hMSCs, fibroblasts and many other cell types^[6]^, but is highly unstable in solution^[6, 7]^. To potentially enhance the regeneration capacity, we set out to covalently couple bFGF to ESP scaffolds.

300PEOT45PBT55 was used to produce 50μm thick ESP scaffolds with 0.99±0.18μm average fiber diameter (Supplementary Fig. 1).The ester bond in the polymer was opened using 0,5M NaOH to expose carboxyl groups on the surface of the scaffold. EDC-NHS chemistry was used to covalently couple the free amine groups of proteins to the surface of the scaffold. As a model protein, FITC labeled bovine serum albumin (BSA) was coupled to the ESP scaffolds. A ~27 fold (26.4±0.8x, p<0.001) higher fluorescent signal was observed when BSA was added after EDC-NHS, than when BSA was added after water control (Fig. 1a). After washing with SDS, to wash away non-covalently bound BSA, the fluorescent signal was 39.8±14.5x higher (p<0.001) in the EDC-NHS group compared to BSA only. Together, this strongly suggests that covalent coupling of BSA was achieved.

**Figure 1.**
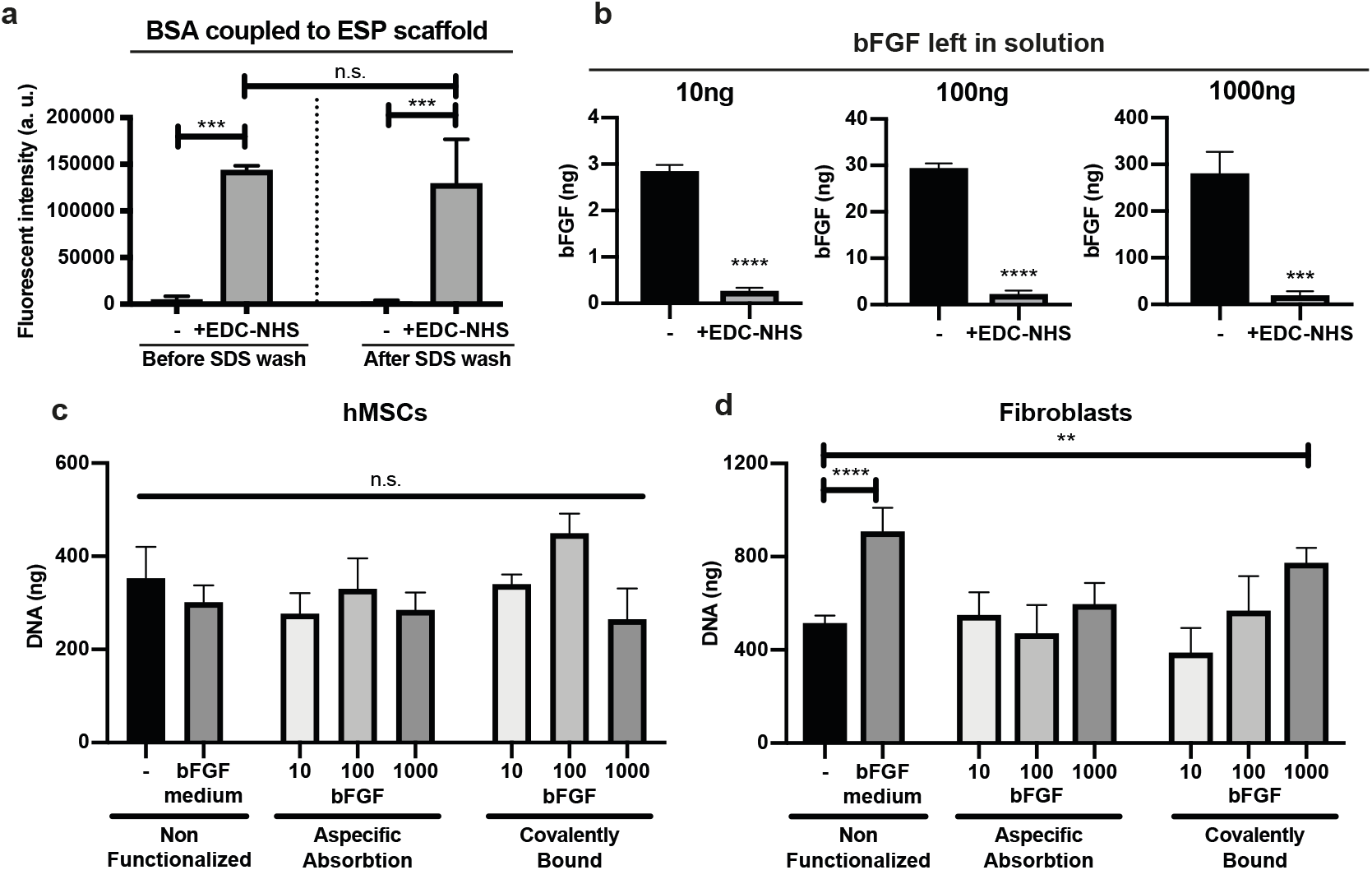
Functional coupling of bFGF to ESP scaffold. **a,** Fluorescent model protein BSA coupled to ESP scaffold using EDC-NHS, or water (-). Right bars are the same scaffolds after overnight wash with 1% (w/v) SDS in water. n=3 scaffolds per condition. **b,** Measurement of bFGF left in solution by ELISA after functionalization of 10, 100 or 1000ng bFGF per scaffold, using EDC-NHS+bFGF, or water+bFGF(-). n=3 scaffolds per condition. **c, d,** DNA quantification of hMSCs **(c)** or human dermal fibroblasts **(d)** cultured on ESP scaffolds functionalized with 10, 100 or 1000ng bFGF per scaffold using bFGF+EDC-NHS (covalently bound, right 3 bars), bFGF+water (aspecific absorbtion, middle 3 bars), or non-functionalized scaffolds (left two bars). Cells were cultured in basic medium, or in medium supplemented with 10ng/ml bFGF (bFGF medium condition). n=3 scaffolds per condition for **(c)**, and n=5 scaffolds per condition for **(d)**. **a, c, d,** One-way ANOVA with Tukey’s post-hoc test. **b,** Student’s t-test. **a-d,** n.s. p > 0,05; ** p<0,01; *** p<0,001; **** p<0,0001. Error bars indicate mean±SD.

Next, bFGF was coupled to ESP scaffolds, using the same strategy. As opposed to bFGF in solution, cell response to covalently coupled bFGF has not been widely studied. In an attempt to find the right concentration range, we coupled three different amounts of bFGF to ESP scaffolds. bFGF left over in solution after coupling was measured by ELISA (Fig. 1b). Without the addition of EDC-NHS, around 70% of the bFGF adhered aspecifically to the scaffolds in all three concentrations. With the addition of EDC-NHS, around 98% of bFGF was bound to the scaffolds, aspecifically or covalently. Before cell culture, scaffolds were thoroughly washed in water and PBS, to wash away most of the aspecifically bound bFGF.

To test whether the bound bFGF was still functional, proliferation of hMSCs cultured on the ESP scaffolds was assessed after 7 days (Fig. 1c). Interestingly, the hMSCs did not respond to either bFGF bound to the ESP scaffold, or bFGF in solution. In 2D tissue culture plastic, hMSCs did increase proliferation over 7 days in response to bFGF in solution, displaying 45±11% more DNA, demonstrating that the ESP scaffold environment influenced the hMSC’s response to bFGF (Supplementary Fig. 2).

Fibroblasts are particularly well studied for their increase in proliferation in response to bFGF. To test whether this lack of response to bFGF when cultured on ESP scaffolds was specific to hMSCs, human dermal fibroblasts were cultured for 7 days on the ESP scaffolds. On non-functionalized scaffolds, 76.5±19.6% more (p<0.0001) DNA was found after 7 days of culture in the presence of bFGF in the medium. On the 1000ng covalently coupled bFGF scaffolds, 50.2±12.5% more (p<0.01) DNA was found compared to non-functionalized scaffolds, showing that the covalently bound bFGF was still functional.

Heparin is known to bind and stabilize bFGF and increase efficacy^[47]^. To covalently couple heparin to ESP scaffolds, PEG-NH2 was incorporated into the electrospinning polymer solution to introduce amino groups on the surface of the ESP scaffold. The carboxyl groups of heparin were then bound to the ESP scaffolds by EDC-NHS chemistry (Supplementary Fig. 3a). bFGF was then bound to the heparin-functionalized scaffolds by overnight incubation. As the heparin interfered with the bFGF ELISA (data not shown), the amount of absorbed bFGF could not be measured. After 7 days of culture, no differences were observed between hMSCs cultured on heparin+bFGF scaffolds and the heparin only- or non-functionalized scaffolds Supplementary Fig. 3b). This further demonstrates that hMSCs don’t respond to bFGF on ESP scaffolds, also not when bound to heparin.

Together, these results show that the covalently coupled bFGF was still functional, and that hMSCs do not respond to bFGF when cultured on ESP scaffolds, but fibroblasts do.

### Reduced FGFR1 expression on ESP scaffolds in hMSCs, but not fibroblasts

To test why fibroblasts did, but hMSCs did not, respond to bFGF when cultured on such microfibrillar susbtrates, we analyzed FGF receptor 1 (FGFR1) expression of hMSCs and fibroblasts cultured on TCP, ESP scaffolds, and on 2D films made up of the same material as the ESP scaffolds. Interestingly, when cultured on ESP scaffolds, hMSCs expressed 86.5±5.3% less (p<0.01) FGFR1 than when cultured on TCP (Fig. 2a). On films, hMSCs displayed 66.7±6.6% less (p<0.01) FGFR1 expression than on TCP, showing that part of the reduction of FGFR1 expression on ESP scaffolds comes from the material properties. However, on ESP scaffolds the FGFR1 expression was still 59.5±15.8% lower (p<0.05) than on films, showing that regardless of material properties, the microfibrillar environment influenced FGFR1 expression.

**Figure 2.**
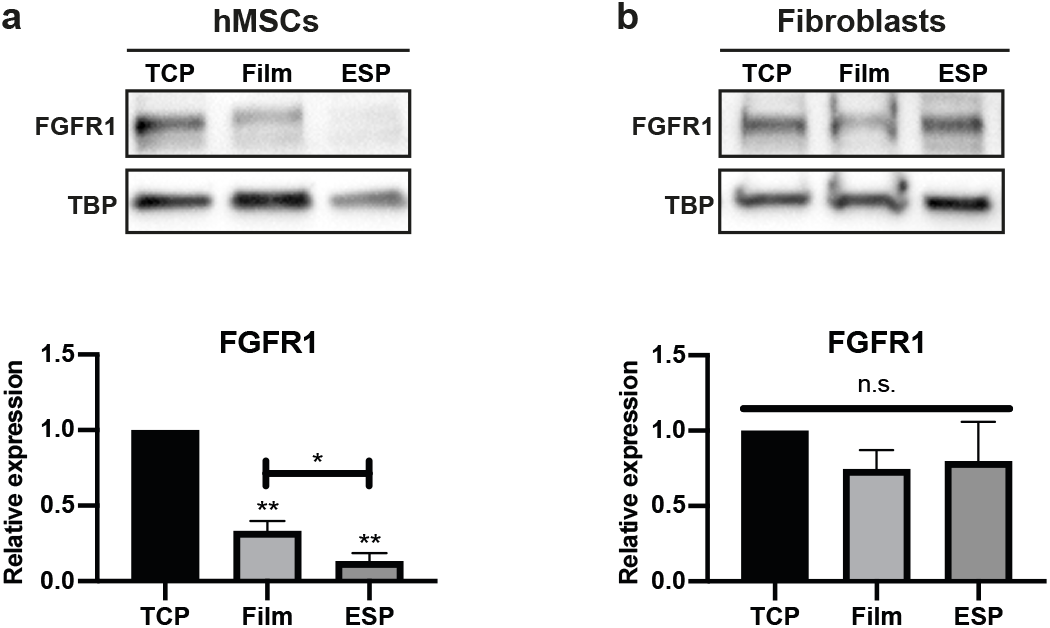
FGFR1 expression of hMSCs and fibroblasts on TCP, films and ESP scaffolds. **a, b,** Western blot of FRFR1 and TBP (as loading control) of hMSCs **(a)** or human dermal fibroblasts **(b)** on TCP, films or ESP scaffolds. Graphs depict quantification of western blots of FGFR1/TBP from 4 **(a)**, or 3 **(b)** independent experiments, normalized to TCP. **a, b,** Repeated measures ANOVA with Tukey’s post-hoc test. Stars above bars indicate significance compared to TCP. n.s. p>0,05; * p<0,05; ** p<0,01. Error pars indicate mean±SD.

Fibroblasts, however, did not display a difference in FGFR1 expression between the different culture substrates (Fig. 2b). The reduced FGFR1 expression of hMSCs on ESP scaffolds, and the high FGFR1 expression of fibroblasts on ESP scaffolds, potentially explains the difference in bFGF response of hMSCs and fibroblasts on ESP scaffolds.

### hMSCs, but not fibroblasts, display fewer focal adhesions on ESP scaffolds

To understand why hMSCs, but not fibroblasts, reduced FGFR1 expression on ESP scaffolds, we investigated the difference in adhesion to the different substrates in hMSCs and fibroblasts by looking at focal adhesions. The expression of zyxin, an important focal adhesion protein, was reduced in both hMSCs and fibroblasts, respectively by 65.6±7.0% (p<0.01) and 79.4±10.9% (p<0.05) compared to TCP (Fig. 3a, b). Also paxillin expression, another well studied focal adhesion protein, was significantly reduced in both hMSCs and fibroblasts on ESP scaffolds, compared to TCP; respectively 73.2±5.2% (p<0.01) and 65±7.9% (p<0.05) (Supplementary Fig. 4a, b). On films, hMSCs also displayed reduced zyxin and paxillin expression, respectively 63.1±16.7% and 41.4±11.4% compared to TCP. Fibroblasts did not show a significant difference in zyxin or paxillin expression on films, compared to TCP.

**Figure 3.**
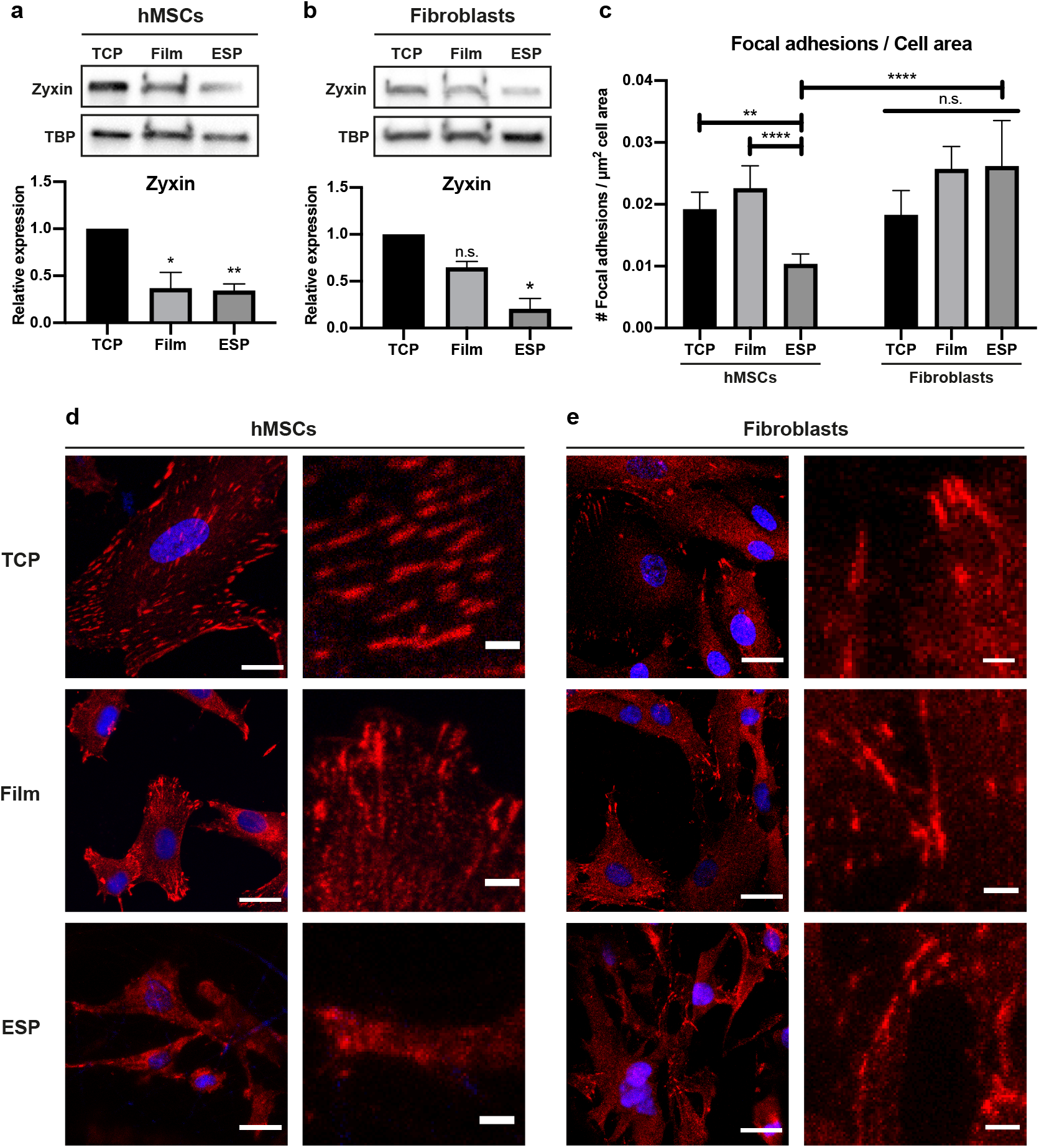
Zyxin expression and focal adhesion analysis of hMSCs and fibroblasts on TCP, films and ESP scaffolds. **a, b,** Western blot of zyxin and TBP (as loading control) of hMSCs **(a)** or human dermal fibroblasts **(b)** on TCP, films or ESP scaffolds. Graphs depict quantification of western blots of zyxin/TBP from 4 **(a)**, or 3 **(b)** independent experiments, normalized to TCP. Stars indicate significance compared to TCP. Repeated measured ANOVA with post-hoc test. Error bars indicate mean±SD. **c,** quantification of number of zyxin positive focal adhesions per μm2 cell area of hMSCs or human dermal fibroblasts grown on TCP, films or ESP scaffolds. n=17-27 cells, quantified in 5-10 different images from biological triplicates. One-way ANOVA with post-hoc test. Error bars indicate mean±95% CI. **a, b, c,** n.s. p>0,05; * p<0,05; ** p<0,01; **** p<0,0001. **d, e,** Representative images of hMSCs **(d)** or human dermal fibroblasts **(e)** stained for zyxin (red) and nuclei (blue). Right panels represent a 5x magnification of the respective left panel. Scalebars represent 25μm (left panels) and 4μm (right panels).

When looking at the formation of zyxin positive focal adhesions, a reduction of 46.0±18.4% (p<0.01) of focal adhesions per cell area was observed when hMSCs were cultured on ESP scaffolds, compared to TCP (Fig. 3c, d). Also when compared to films, hMSCs on ESP scaffolds displayed 54.1±15.7% (p<0.0001) less zyxin positive focal adhesions per cell area. Interestingly, no significant difference was found between fibroblasts cultured on the different substrates (Fig. 3c, e). Indeed, when compared to fibroblasts grown on ESP scaffolds, hMSCs on ESP scaffolds displayed 60.4±13.5% (p<0.0001) fewer focal adhesions per cell area. The same trend was observed for paxillin positive focal adhesions, where hMSCs displayed far fewer paxillin positive focal adhesions on ESP scaffolds than on films or TCP, while fibroblasts contained many paxillin positive focal adhesions on all three substrates (Supplementary Fig. 4c, d).

These results demonstrate that the microfibrillar environment of ESP scaffolds changes focal adhesion formation in hMSCs, but not in fibroblasts. This shows that hMSCs adhere differently to the ESP scaffolds than fibroblasts, potentially explaining the difference in FGFR1 expression.

### FGFR1 is not regulated through zyxin or paxillin

As the lower FGFR1 expression correlated with fewer focal adhesions of hMSCs on ESP scaffolds, we knocked down paxillin and zyxin. Interestingly, neither paxillin nor zyxin depletion resulted in a change in FGFR1 expression, demonstrating that the differential expression of these proteins by hMSCs on ESP scaffolds is not the reason for the reduced FGFR1 expression (Figure 4a-b).

**Figure 4.**
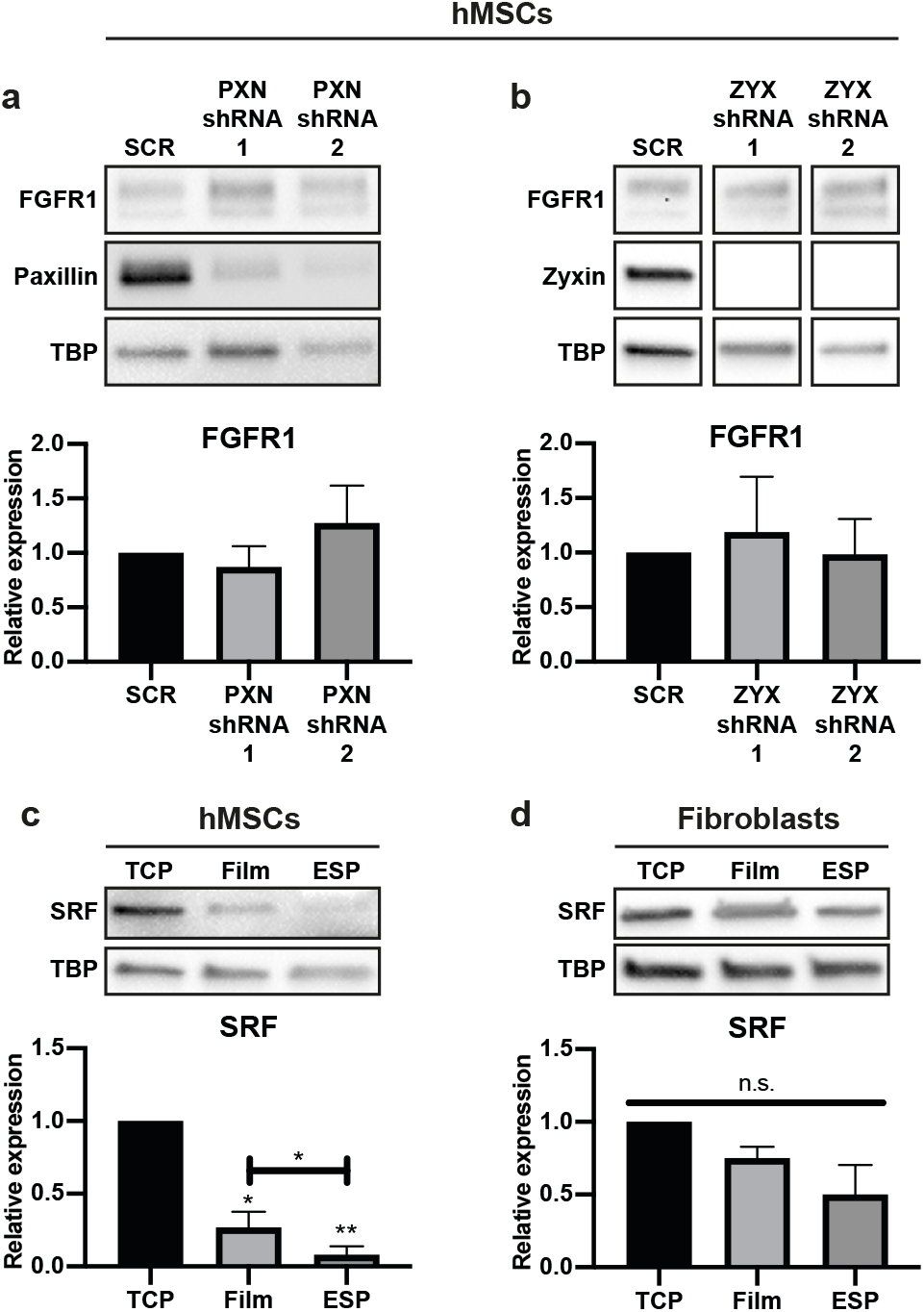
No role of paxillin or zyxin in regulation of FGFR1. **a, b,** Western blot of FGFR1, paxillin, zyxin and TBP (as loading control) of hMSCs transduced with PXN-shRNA **(a)** or ZYX-shRNA **(b)** cultured on TCP. Graphs depict quantification of western blots of FGFR1/TBP from 4 biological replica’s, normalized to TCP. **c, d,** Western blot of SRF and TBP (as loading control) of hMSCs **(c)** or human dermal fibroblasts **(d)** cultured on TCP, films or ESP scaffolds. Graphs depict quantification of western blots of SRF/TBP from 4 **(c)** or 3 **(d)** independent experiments, normalized to TCP. Stars above bars indicate significance compared to TCP. **a, b, c, d,** Repeated measures ANOVA with Tukey’s post-hoc test. n.s. p>0,05; * p<0,05; **** p<0,0001. Error bars indicate mean±SD.

Even though focal adhesions didn’t influence the FGFR1 expression, the reduction in focal adhesions of hMSCs on ESP suggests a difference in mechanosensitive signaling. An important mechanosensitive pathway is the MRTF/SRF pathway. MRTF translocates to the nucleus when actin is incorporated into actin filaments and globular actin is low, where it activates SRF to transcribe target genes. To investigate this pathway, we looked at the expression of SRF. Indeed, compared to TCP, the SRF expression was 73.1±10.7% (p<0.05) lower on films and 91.9±5.9% (p<0.01) lower on ESP scaffolds in hMSCs. Compared to films, SRF expression was 69.7±21.8% (p<0.05) lower on ESP scaffolds (Fig. 4c). For fibroblasts, expression of SRF was 24.9±79% (p>0.05) and 50±20.3% (p>0.05) lower on films and ESP respectively, but this difference was not statistically significant (Fig. 4d). To further investigate the MRTF/SRF pathway, we looked at the localization of MRTF-A in hMSCs and fibroblasts, on TCP, films and ESP scaffolds. MRTF-A was mainly located in the nucleus in fibroblasts, regardless of the culture substrate (Supplementary Fig. 5). In hMSCs, however, MRTF-A was located in the nucleus when cultured on TCP, but mainly in the cytoplasm when cultured on films. Surprisingly, hMSCs on ESP scaffolds also showed nuclear localization of MRTF-A. Together with the SRF expression, these results suggest activity of the MRTF/SRF pathway in fibroblasts on all substrates and of hMSCs on TCP, but not of hMSCs on films or ESP scaffolds. The activity of the MRTF/SRF pathway correlates with the reduced FGFR1 expression of hMSCs on films or ESP scaffolds.

### Actin-myosin and MRTF/SRF pathway regulate FGFR1 expression

To investigate the role of the MRTF/SRF pathway in the regulation of FGFR1 in hMSCs, we inhibited the pathway using CCG203971^[48, 49]^. Indeed, in both hMSCs and fibroblasts, inhibition of the MRTF/SRF pathway reduced FGFR1 expression by 59.6±7.0% (p<0,01) and 61.5±2.9% (p<0.01), respectively (Fig 5a, c). This shows that MRTF/SRF directly or indirectly regulates FGFR1 expression in both hMSCs and fibroblasts. We observed a strong decrease in SRF expression in hMSCs on ESP scaffolds (Fig. 4c), strongly suggesting that the reduced FGFR1 expression of hMSCs on ESP scaffolds is due to a decrease in SRF expression. Fibroblasts maintained a high expression of SRF on ESP scaffolds (Fig. 4d), explaining the high expression of FGFR1 on ESP scaffolds.

**Figure 5.**
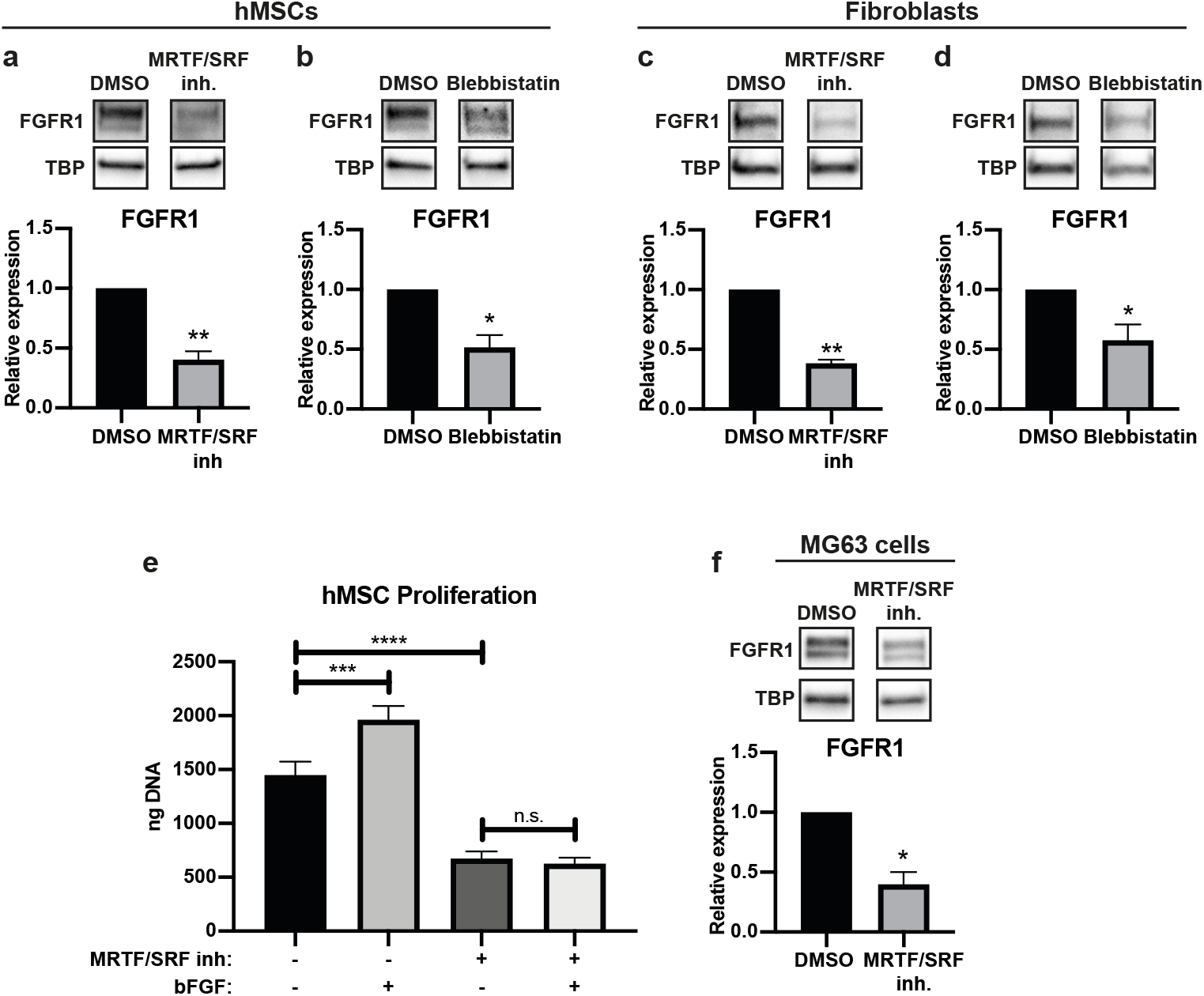
Actin-myosin and MRTF/SRF inhibitors change FGFR1 expression in hMSCs, fibroblasts and MG63 cells. **a, b, c, d,** Western blot of FGFR1 and TBP (as loading control) of hMSCs **(a, b)** or human dermal fibroblasts **(c, d)**, cultured on TCP and treated with MRTF/SRF inhibitor CCG203971 **(a, c)** or blebbistatin **(b, d)**. Graphs depict quantification of western blots of FGFR1/TBP from 3 biological replica’s, normalized to TCP. **e,** DNA quantification of hMSCs cultured for 7 days on TCP in the presence of MRTF/SRF inhibitor and/or 10ng/ml bFGF. n=3 for each condition. One-way ANOVA with Tukey’s post-hoc test. n.s. p>0,05; *** p<0,001; **** p<0,0001; Error bars indicate mean±SD. **f,** Western blot of FGFR1 and TBP (as loading control) of MG63, an osteosarcoma cell-line cultured on TCP and treated with MRTF/SRF inhibitor CCG203971. Graph depicts quantification of western blots of FGFR1/TBP from 3 biological replicas, normalized to TCP. **a, b, c, d, f,** ratio paired t-test. * p<0,05; ** p<0,01. Error bars indicate mean±SD.

When most actin monomers are assembled into filaments and globular actin is low, the MRTF/SRF pathway is activated. To determine the role of the actin cytoskeleton in the regulation of FGFR1, we treated hMSCs and fibroblasts with blebbistatin, an inhibitor of actin-myosin interaction. Expression of FGFR1 reduced 48.4±10.2% (p<0.05) in hMSCs and 42.4±13.3% in fibroblasts (Fig. 5b, d). Together, this demonstrates that FGFR1 is regulated by the actin cytoskeleton, directly or indirectly through the MRTF/SRF pathway.

Another important mechanosensitive co-transcription factor is Yes activated protein 1 (YAP), entering the nucleus when a cell experiences high cellular tension^[33]^. To investigate if YAP plays a role in FGFR1 regulation, we knocked down YAP in hMSCs. No difference was observed in FGFR1 expression between YAP-knock down and control-shRNA groups (Supplementary Fig. 6), demonstrating that YAP does not play a role in FGFR1 regulation in hMSCs.

To further investigate the link between the MRTF/SRF pathway and the FGF pathway, we investigated the response to bFGF of hMSCs cultured with MRTF/SRF inhibitor. After 7 days of culture on TCP in the presence of bFGF and/or MRTF/SRF inhibitor, total DNA was analyzed. As expected, 35.5±8.9% more DNA was found when bFGF was added to the medium, compared to basic medium (Fig. 5e). In the presence of MRTF/SRF inhibitor, 53.4±4.6% less DNA was found than in basic medium. Interestingly, in the presence of MRTF/SRF inhibitor, hMSCs did not increase proliferation when bFGF was added. This shows that the MRTF/SRF pathway regulates the response to bFGF, in confirmation with the reduced FGFR1 expression.

Aberrant FGFR regulation in cancer cells has been linked to metastasis, tumor progression and a worse diagnosis. To test whether the MRTF/SRF pathway is also responsible for FGFR1 regulation in cancer cells, we treated the osteosarcoma cell line MG63 with the MRTF/SRF inhibitor. Similar to hMSCs and fibroblasts, FGFR1 expression was reduced by 60.2±10.3% (p<0.05) when MRTF/SRF was inhibited (Fig. 5f). MRTF/SRF inhibition decreased FGFR1 expression in 3 different human cell types, suggesting that the MRTF/SRF pathway is a univocal regulator of the FGFR1 pathway.

## Discussion

Here, we have functionalized 300PEOT45PBT55 ESP scaffolds by coupling bFGF to the surface. The covalent binding of bFGF to ESP scaffolds made of other polymers has been shown before to retain the growth factor bioactivity^[10, 50]^. Similarly, the covalently coupled bFGF was still active on our ESP scaffolds, and could be used as a method to increase cell proliferation rate on ESP scaffolds. This could be useful for *in vivo* approaches, but it can also be used as a cell culture substrate *in vitro*. bFGF is highly unstable in solution and covalent binding to a surface has been shown to increase its stability^[10]^.

Unlike fibroblasts, hMSCs did not increase proliferation in response to bFGF (in solution or covalently bound) on ESP scaffolds. We found that this was due to reduced SRF expression, which caused decreased FGFR1 expression. SRF expression is known to be regulated by itself through a positive feedback loop^[51]^. The observed difference in SRF expression between TCP, films and ESP scaffolds highlight the difference in SRF activity on the different substrates. The positive feedback loop can increase the differences in SRF expression, but the origin of the initial difference in SRF expression remains unclear. MRTF-A was located in the nucleus of fibroblasts on all substrates, and of hMSCs on TCP, strongly suggesting together with high SRF activity that the MRTF/SRF pathway was active. hMSCs on films did not show nuclear localization of MRTF-A, which together with the low SRF expression suggests that the pathway is inactive, explaining the low FGFR1 expression. On ESP, however, MRTF-A was located in the nucleus. Even though MRTF-A was located in the nuclei, the low SRF expression of hMSCs on ESP scaffolds could prevent active transcription of the FGFR1 gene, or of genes that (indirectly) regulate FGFR1.

Through actin-myosin inhibition by blebbistatin, we found that FGFR1 expression is reduced with less actin-myosin tension. The MRTF/SRF pathway is dependent on the actin cytoskeleton, but also plays a role in shaping the actin network^[52]^. Whether the effect of actin-myosin inhibition went through MRTF/SRF, or vise-versa, we did not investigate. It is possible that no clear cause and effect between these two players exists, because there is a positive feedback look between the two. MRTF/SRF activity increases stress fiber formation, thereby also increasing MRTF nuclear localization and increasing MRTF/SRF activity^[52]^. hMSCs grown on ESP scaffolds displayed fewer focal adhesions than on films or TCP. In contrast, fibroblasts formed similar numbers of focal adhesions per cell area on ESP scaffold as on films or TCP. On TCP and films, the number of zyxin positive focal adhesions was the same between hMSCs and fibroblasts. The reason for the difference between fibroblasts and hMSCs was not investigated here. Different cell types exhibit different cell spreading and traction forces in response to different substrate stiffnesses^[53–55]^. Indeed, the optimal stiffness for differentiation and proliferation defer per cell type^[56, 57]^. We have previously shown that hMSCs experience the ESP scaffolds used here as a soft substrate, demonstrated by fewer focal adhesions, less lamin A/C and less YAP nuclear translocation^[58]^. The difference in focal adhesion formation between hMSCs and fibroblasts observed here potentially derives from a different response to matrix stiffness. Perhaps fibroblasts are able to form focal adhesions on softer substrates than hMSCs. Side by side comparison of hMSCs and fibroblasts on different stiffnesses has not yet been reported, but could shed light on the differences observed here.

Knockdown of either zyxin or paxillin did not affect FGFR1 expression. In contrast to paxillin, zyxin knockdown is known to diminish stress fibers^[59–61]^. While actin-myosin inhibition by blebbistatin did decrease FGFR1 expression, zyxin knockdown did not. Although fewer than normally, zyxin knockdown cells still form focal adhesions^[62]^. Our data suggests that the actin-myosin tension between these focal adhesions is still sufficient to maintain a higher FGFR1 expression, as full inhibition of actin-myosin by blebbistatin reduced FGFR1.

The regulation of FGFR1 expression is poorly studied. YAP knockdown has been shown to decrease FGFR1 expression in lung cancer cells^[63]^ and neurospheres^[64]^. Also, integrin α6 has been shown to regulate FGFR1^[64]^. We found that YAP knockdown didn’t alter FGFR1 expression in hMSCs, suggesting a different role for YAP in different cell types. YAP and integrin α6 regulation of FGFR1 does, however, hint at the mechanosensitive regulation of FGFR1, in accordance with what we’ve shown here. Other mechanisms of FGFR1 regulation include regulation by Pdx-1^[65,66]^ and ZEB1^[64]^. The regulation of FGFR1 by MRTF/SRF and actin-myosin tension presented here adds to the sparse literature on FGFR1 regulation. These novel findings can give insight in tumor development, as aberrant FGFR1 regulation is important in a wide variety of cancers^[12, 17]^. FGFR inhibitors are already being used in clinical trials as novel anti-cancer drugs^[18–23]^. Our study opens up new potential targets for FGFR1 regulation in cancer cells. Also, as an important regulator of proliferation in hMSCs^[27]^ and other cell types^[67, 68]^, this can have implications for scaffold designs. We show here that the scaffold design itself, as well as material properties, can influence the FGFR1 expression. Optimizing scaffold design to influence MRTF/SRF activity and FGFR1 expression could be crucial for certain tissue regeneration applications.

## Conclusion

ESP scaffolds were successfully functionalized with bFGF, which increased the proliferation of fibroblasts, but not hMSCs. hMSCs responded to bFGF on TCP, but reduced FGFR1 expression on ESP scaffolds, explaining their irresponsiveness to bFGF on ESP scaffolds. Fibroblasts maintained a high expression of FGFR1 on ESP scaffolds, explaining the difference in bFGF responsiveness between hMSCs and fibroblasts. hMSCs, but not fibroblasts, displayed fewer focal adhesions and expressed less SRF on ESP scaffolds than on TCP or 2D film controls. In hMSCs and fibroblasts, the inhibition of actin-myosin interaction and MRTF/SRF activity decreased FGFR1 expression. In osteosarcoma MG63 cells, MRTF/SRF inhibition also led to decreased FGFR1 expression. Together, our data shows that hMSCs become irresponsive to bFGF on ESP scaffolds because of a downregulation of SRF, which leads to a decrease in FGFR1. Fibroblasts maintain a high SRF and FGFR1 expression and remain responsive to bFGF on ESP scaffolds.

## Author contributions

J.Z. and S.R. performed the experiments. J.Z. and L.M. designed the study and wrote the manuscript.

## Methods

### Film and ESP scaffold production

Random block co-polymer of poly(ethylene oxide terephthalate) (PEOT) and poly(butylene terephthalate) (PBT), with 300Da PEO and PEOT/PBT ratio (w/w) of 55/45 (300PEOT55PBT45, acquired from PolyVation) was used to produce films and ESP scaffolds. 300PEOT55/PBT45 granules were melted at 180 °C under slight pressure (~100 kg) in a circular 23mm mold between two silicon wafers (Si-mat, Kaufering, Germany) to produce flat films. Films were punched out using a 22mm punch to fit in a 12well plate.

The electrospinning polymer solution was prepared by dissolving 20% (w/v) 300PEOT55PBT45 in 3:7 1,1,1,3,3,3,-Hexafluoro-2-propanol (HFIP):chloroform, overnight at room temperature under agitation. For heparin functionalization, 2% (w/v) poly(ethylene glycol) (PEG) with 2 NH2 end-groups (Mw: 3350kDa) (PEG-NH2), was added to the polymer solution and mixed for 4 hours before electrospinning.

ESP scaffolds were produced on a slowly rotating (100RPM) 19cm diameter mandrel by electrospinning on a polyester mesh (FinishMat 6691 LL (40 gr/m2), generously provided by Lantor B.V.) with 12mm holes, on top of aluminum foil. The following parameters were maintained: 15cm working distance between needle and rotating mandrel, 1ml/hr flow rate, 23-25 °C and 40% relative humidity, a needle charge between 10-15kV and collector charge between −2 and −5kV. Individual ESP scaffolds were punched out with a diameter of 15mm over the 12mm holes in the polyester mesh and removed from the aluminum foil. This resulted in 15mm ESP scaffolds with a 12mm diameter surface for cell culture and a 1,5mm polyester ring around it to improve handleability. Using this method, up to 100 ESP scaffolds were produced under exactly equal parameters.

Before cell culture, ESP scaffolds and films were sterilized in 70% ethanol for 15min and dried at room temperature until visually dry. The 1,5mm polyester ring was covered with a rubber 15mm outer- and 12mm inner-diameter O-ring (Eriks) to keep the scaffolds from floating in tissue culture well plates.

### Functionalization of ESP scaffolds with BSA or bFGF

Before coupling of bovine serum albumin (BSA)-FITC conjugate (ThermoFisher Scientific) or basic fibroblast growth factor (bFGF) (Neuromics), ethanol sterilized ESP scaffolds were incubated in 0,5M NaOH for 30min at room temperature to open the ester bond of the 300PEOT55PBT45 polymer. Scaffolds were thoroughly washed 5 times with water and then incubated with 4mg/ml N-(3-Dimethylaminopropyl)-N’-ethylcarbodiimide hydrochloride (EDC) (Sigma-Aldrich) and 10mg/ml N-hydroxysuccinimide (NHS) (Sigma-Aldrich) in milliQ water, or in milliQ water only, without EDC-NHS, as negative control, for 30minutes at room temperature on a rocking plate. EDC-NHS solution was removed and 500μl of 1μg/ml BSA or 20, 200, or 2000ng/ml bFGF in water was added to the scaffolds in a 24 well-plate well and incubated overnight at 4 °C on a rocking plate. The following day, BSA-FITC scaffolds were washed 5 times with water and scaffold fluorescence was measured in the fluorescein channel on a Clariostar plate reader (BMG Labtech). For sodium dodecyl sulfate (SDS) wash, 1% (w/v) in water was added to the functionalized scaffolds and incubated under agitation at room temperature overnight. The following day, scaffolds were thoroughly washed 5 times with water and measured on the plate reader as described before.

For bFGF functionalized scaffolds, bFGF solution was harvested to be analyzed by bFGF ELISA, and the scaffolds were washed 5 times with water, once with PBS and once with medium. For the bFGF scaffolds, all solutions were sterilized by filtration through at 0,2μm filter. bFGF was quantified using a bFGF ELISA kit (Abcam), according to manufacturer’s protocol.

### Heparin functionalization of ESP scaffolds

1, 5mg/ml heparin sodium salt from porcine intestinal mucosa (Sigma-Aldrich) was mixed with 4mg/ml EDC and 10mg/ml NHS in water (or water only, without EDC-NHS, as negative control) and directly added to the 300PEOT55PBT45+PEG-NH2 ESP scaffolds and incubated overnight at 4 °C.

To measure bound heparin, scaffolds were washed with 5 times with milliQ water and stained for 30min with alcian blue staining solution (0.1% alcian blue, 10% ethanol, 0.1% acetic acid, 0.03M MgCl_2_ in water (all Sigma-Aldrich)). Scaffolds were washed once with MQ water and incubated for 30min at room temperature in destaining solution (10% ethanol, 0.1% acetic acid, 0.03M MgCl_2_ in water). Scaffolds were washed again once with water and then incubated for 30min in 1% SDS to extract the heparin-bound alcian blue from the scaffolds. The absorbance of this solution was measured in a Clariostar plate reader.

For cell culture, the heparin functionalized scaffolds were washed 5 times with milliQ water and incubated overnight at 4 °C with 500μl 2000ng/ml bFGF. The following day scaffolds were washed 5 times with water, once with PBS and once with medium. All solutions were sterilized by filtration through at 0,2μm filter.

### Cell culture

Human dermal fibroblasts (Lonza) were expanded at 2000 cells/cm^2^ in DMEM+Glutamax medium (Thermo Fisher Scientific) supplemented with 10% (V/V) fetal bovine serum (FBS) (Sigma-Aldrich).) Bone marrow derived hMSCs were isolated by Texas A&M Health Science Center^[69]^. Briefly, aspirated bone marrow was centrifuged to isolate mononuclear cells. The hMSCs were further expanded and tested for differentiation potential. hMSCs were received at passage 1 and were further expanded at 1000 cells/cm^2^ in α-MEM+Glutamax medium (Thermo Fisher Scientific) supplemented with 10% FBS. MG63 cells (ATCC) were expanded at 5000 cells/cm^2^ in DMEM+Glutamax+10% FBS medium. All cells were cultured in 37 °C in 5% CO2 until reaching 70-80% confluency. Cells were trypsinised in 0.05% Trypsin and 0.53 mM EDTA (ThermoFisher Scientific) and hMSCs and fibroblasts were used for experiments at passage 5. MG63 cells were used at passage 90.

Unless otherwise stated, all experiments were harvested at day 7. For scaffold experiments, hMSCs and fibroblasts were cultured at 1000 cells/cm^2^ in TCP and films, and 30.000 cells/ESP scaffold in growth medium with and 100 U/ml penicillin-streptomycin. All other experiments were done in medium without penicillin-stretomycin. In bFGF medium conditions, 10ng/ml bFGF was added to the medium.

For blebbistatin and MRTF/SRF inhibitor experiments, hMSCs and fibroblasts cells were seeded at 1000 cells/cm^2^ on TCP and cultured for 6 days. MG63 cells were seeded at 5000 cells/cm^2^ cultured for 2 days, because of a very high proliferation rate. After the initial culture period in growth medium, 100μM blebbistatin (Sigma-Aldrich) in 0,2% DMSO in growth medium, or 12,2μM MRTF/SRF inhibitor CCG203971 in 0,1% DMSO in growth medium, or respective DMSO control was added to the cells for 24h.

To test the responsiveness of hMSCs to bFGF in the presence of MRTF/SRF inhibitor, hMSCs were seeded at 1000 cells/cm^2^ in TCP and cultured for 7 days in 0,1% DMSO control, 0,1% DMSO + 10ng/ml bFGF, 24,4μM MRTF/SRF in 0,1% DMSO or 24,4μM MRTF/SRF in 0,1% DMSO + 10ng/ml bFGF, all in hMSC growth medium.

### DNA quantification

To lyse cells for DNA quantification, cells were washed 2x with PBS and freeze-thawed dry twice before RLT lysis buffer (Qiagen) was added. ESP scaffolds were removed from the polyester ring after the last PBS wash. Samples were then freeze-thawed 3x in lysis buffer. TCP plates and films were scraped with a cell scraper after the first freeze-thaw in lysis buffer. ESP scaffolds were left in lysis buffer. Samples were diluted 100-400x, depending on expected number of cells per samples, in Tris-EDTA buffer (10 mM Tris-HCl, 1 mM EDTA, pH 7.5) and λ-DNA standard was made in the same final solution (0,25-1% RLT in Tris-EDTA buffer). Pico green assay (ThermoFisher Scientific) was then used to quantify DNA, according to manufacturer’s protocol.

### Protein isolation and western blot

Protein was isolated in a custom lysis buffer to allow for the detection of membrane proteins with western blot. Other buffers, such as RIPA buffer, were tried for FGFR1 western blot without success (data not shown). The buffer consisted of 150 mM NaCl, 0.5% sodium deoxycholate, 1% SDS, 1% NP-40 and 50 mM Tris-HCl in water, set to pH 7.4. The buffer was supplemented with cOmplete™ Mini EDTA-free Protease Inhibitor Cocktail (Sigma-Aldrich). Samples were washed in cold PBS twice before lysis. ESP scaffolds were removed from the supporting polyester ring. To get sufficient proteins, 6–12 films or 15–20 ESP scaffolds were combined in 300-400μl for a single protein isolate. Experiments were repeated 3 or 4 times to obtain sufficient replicates. 6 or 10 cm dishes were used for TCP samples. TCP and film conditions were scraped in lysis buffer with cell scrapers. ESP scaffolds were submerged in lysis buffer and incubated for around 30min in lysis buffer because the scaffolds were removed from the protein isolate. Samples were not spun down to maintain potentially non-dissolved membrane proteins in solution.

Pierce BCA protein assay kit (Thermo Fisher Scientific) was used to quantify total protein concentration. 20μg protein was incubated in 10% 2-Mercaptoethanol (Sigma-Aldrich) in laemmli loading buffer (Bio-Rad) for 37 °C for 20min for FGFR1 western blots and at 95 °C for 5min for all other western blots. Samples were loaded into 4–15% polyacrylamide gels (Bio-Rad) and blotted to 0,45μm PVDF membranes (Bio-Rad) using semi-dry transfer. Membranes were blocked in 5% (w/v) fat free milk (Bio-Rad) in TBS + 0,05% (v/v) tween-20 (Sigma-Aldrich) for 1 hour, except for SRF western blots, which had to be blocked in 2% (w/v) BSA (VWR) + 0,05% tween-20 in PBS to work (data not shown). Primary antibodies were incubated in their respective blocking buffer overnight at 4 °C. All antibodies were ordered from Abcam: FGFR1: ab76464 1/500; Paxillin: ab32084 1/1000; Zyxin: ab58210 1/1000; YAP1: ab52771 1/1000; SRF: ab53147 1/250; TBP: ab51841 1/1000. Blots were incubated the following day with 0,33μg/ml Goat-anti-rabbit or -mouse HRP (Bio-Rad) in blocking buffer for 1h at room temperature. To visualize the protein bands, blots were incubated with Clarity Western ECL (Bio-Rad) for 1-5min right before imaging.

### Immunofluorescence and imaging

Cells were fixed with 3,6% (v/v) paraformaldehyde (Sigma-Aldrich) in PBS for 20min at room temperature. To block and permeabilize, fixed cells were incubated in 2% (w/v) BSA+0,1% (v/v) triton X (VWR) in PBS. Zyxin or paxillin (Abcam, ab58210 and ab32084, respectively, both 1/1000) were incubated in 2% (w/v) BSA+0,05% (v/v) tween-20 in PBS overnight at 4 °C. The following day, 1/1000 Goat-anti-mouse Alexa Fluor 568 or Goat-anti-rabbit Alexa Fluor 488 was incubated overnight at 4 °C in 2% (w/v) BSA+0,05% (v/v) tween-20 in PBS. The next day, samples were stained with DAPI (Sigma-Aldrich, 0,14μg/ml in PBS+0,05% (v/v) tween-20) to stain nuclei. Images were taken on a confocal microscope.

Focal adhesions were quantified manually by counting the number of focal adhesions per cell using Fiji. Between 17 and 27 cells were counted per condition, from 5-10 different images from biological triplicates. Cell area was measured by manually outlining the cells and measuring surface area using Fiji. The number of focal adhesions was normalized to the cell area.

### Lentiviral production and transduction

To produce lentiviral particles, human embryonic kidney 293FT (HEK) cells were seeded at 60.000 cells/cm^2^ in DMEM+Glutamax+10% FBS. Cells were transfected with pMDLg pRRE, pMD2.G, pRSV Rev (Addgene) and one of the pLKO.1 shRNA plasmids using 5:1 lipofectamine 2000 (Thermo Fisher Scientific):DNA (v/w) 24h after seeding. The following TRC pLKO.1 constructs (Dharmacon) were used: ZYX: TRCN0000074204 and TRCN0000074205; PXN: TRCN0000123134 and TRCN0000123136; YAP1: TRCN0000107265 and TRCN0000107266; and non-targeting shRNA control (RHS6848). Medium was changed 16h post-transfection to hMSC growth medium. Lentivirus was harvested and filtered through a 0.45μm filter 24h and 48h after the change to hMSC growth medium.

24h after thawing at 1000 cells/cm^2^, hMSCs were transduced with the lentiviral medium for 16 hours. Medium was replaced with growth medium the following day. 48-72 hours post transduction, medium was replaced with growth medium + 2μg/ml puromycin for 72 hours. A total of 9-10 days after thawing, hMSCs were passaged and seeded at 1000 cells/cm^2^ on TCP for 7 days in growth medium before protein harvest.

### Statistics

The statistical tests and number of biological replicates and/or experiments are stated in the figure subtexts. Each experiment used at least 3 biological replicas. Cells selected for quantification of focal adhesions were selected randomly. Films and electrospun scaffolds were also randomly assigned to different experimental groups. Shapiro-Wilk test was used to test for normal distribution of each experimental group before further statistical analysis. To test for significance of absolute differences in experiments with multiple comparisons between groups, a One-way ANOVA with Tukey’s post hoc was performed. For relative differences between multiple experimental groups, log values were used for repeated measures ANOVA, with Tukey’s post-hoc test. For experiments with a single comparison, two-tailed student’s t-test was used for absolute differences, and ratio-paired t-test for relative differences. Significance was set at p<0.05. Statistical analysis was done using Graphpad Prism 8.

## Acknowledgements

We would like to thank Matt Baker and Paul Wieringa for the valuable discussions about the functionalization strategy of the electrospun scaffolds. We are grateful to the European Research Council starting grant “Cell Hybridge” for financial support under the Horizon2020 framework program (Grant #637308). Some of the materials used in this work were provided by the Texas A&M Health Science Center College of Medicine Institute for Regenerative Medicine at Scott & White through a grant from NCRR of the NIH (Grant #P40RR017447).

## Supplementary Figures

**Supplementary figure 1.**
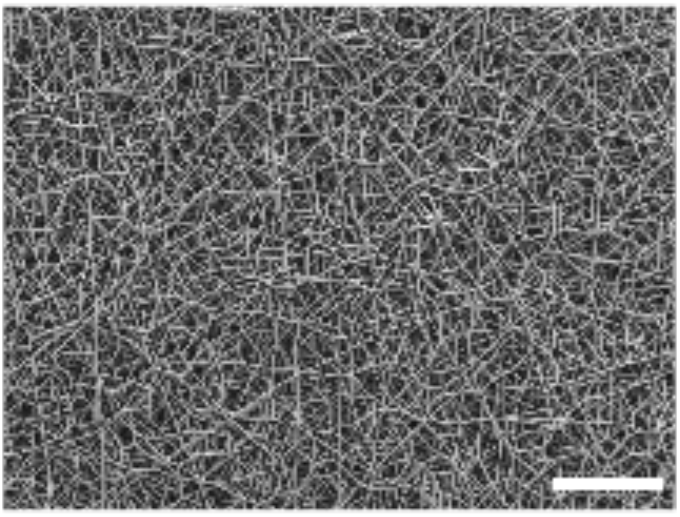
Overview of ESP scaffold. Scalebar 100μm.

**Supplementary figure 2.**
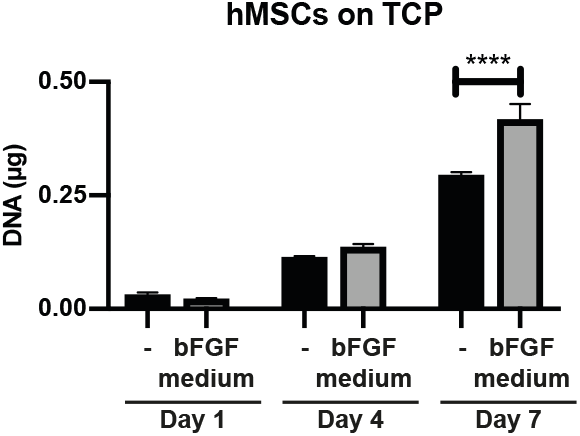
Increased proliferation of hMSCs in response to bFGF. DNA quantification of hMSCs cultured on TCP in basic medium (-) or basic medium + 10ng/ml bFGF harvested on day 1, 4 or 7. n=3 for each condition. One-way ANOVA with Tukey’s post-hoc test. **** p<0,0001.

**Supplementary figure 3.**
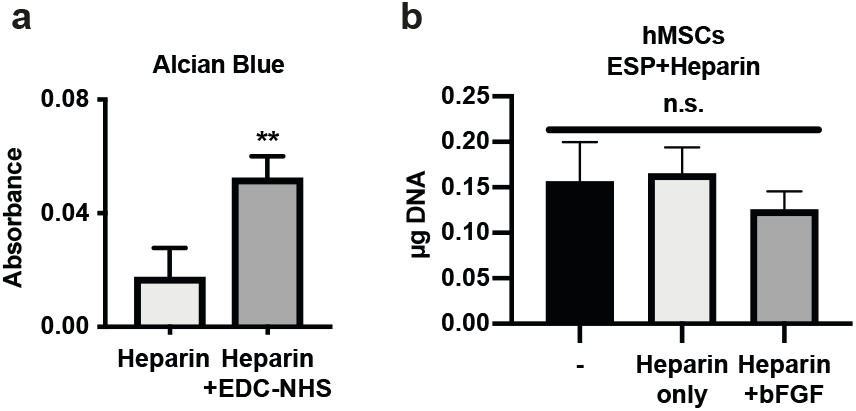
No effect heparin bound bFGF on hMSC proliferation. **a,** Alcian blue analysis of heparin bound to ESP scaffolds with incorporated PEG-NH2 by EDC-NHS chemistry, or aspecifically absorbed heparin, without EDC-NHS. Student’s t-test. ** p<0,01. **b,** DNA quantification of hMSCs cultured for 7 days on unfunctionalized scaffolds (-), or scaffolds functionalized with heparin and with or without absorbed bFGF. One-way ANOVA with Tukey’s post-hoc test. n.s. p>0,05. **a, b,** n=3 for each condition. Error bars indicate mean±SD.

**Supplementary figure 4.**
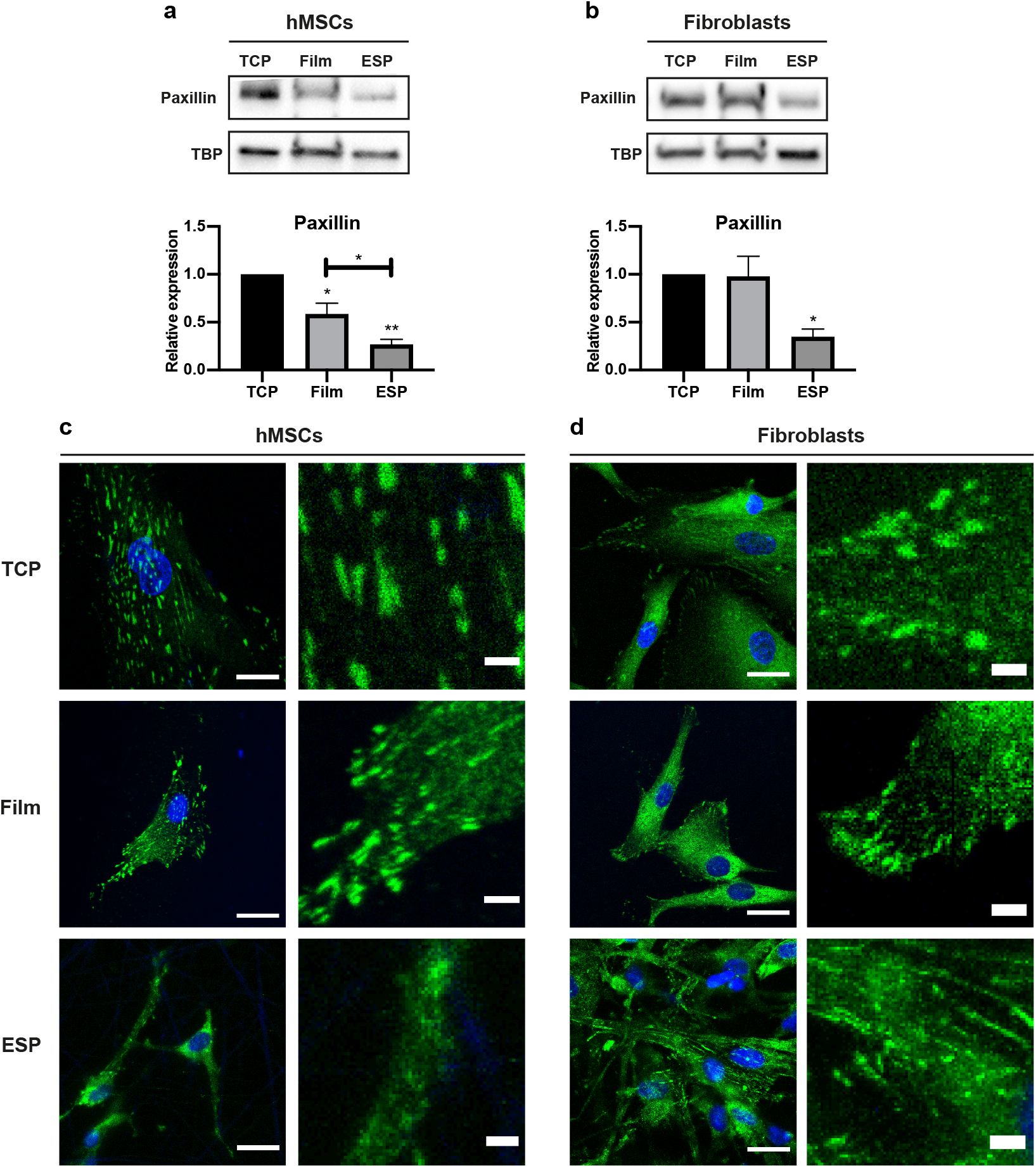
Reduced paxillin expression on ESP scaffolds. **a, b,** Western blot of paxillin and TBP (as loading control) of hMSCs **(a)** or human dermal fibroblasts **(b)** on TCP, films or ESP scaffolds. Graphs depict quantification of western blots of paxillin/TBP from 4 **(a)**, or 3 **(b)** independent experiments, normalized to TCP. Stars above bars indicate significance compared to TCP. Repeated measures ANOVA with Tukey’s post-hoc test. * p<0,05; ** p<0,01. Error bars indicate mean±SD. **c, d,** Representative images of hMSCs **(c)** or human dermal fibroblasts **(d)** stained for paxillin (green) and nuclei (blue). Right panels represent a 5x magnification of the respective left panel. Scalebars represent 25μm (left panels) and 4μm (right panels).

**Supplementary figure 5.**
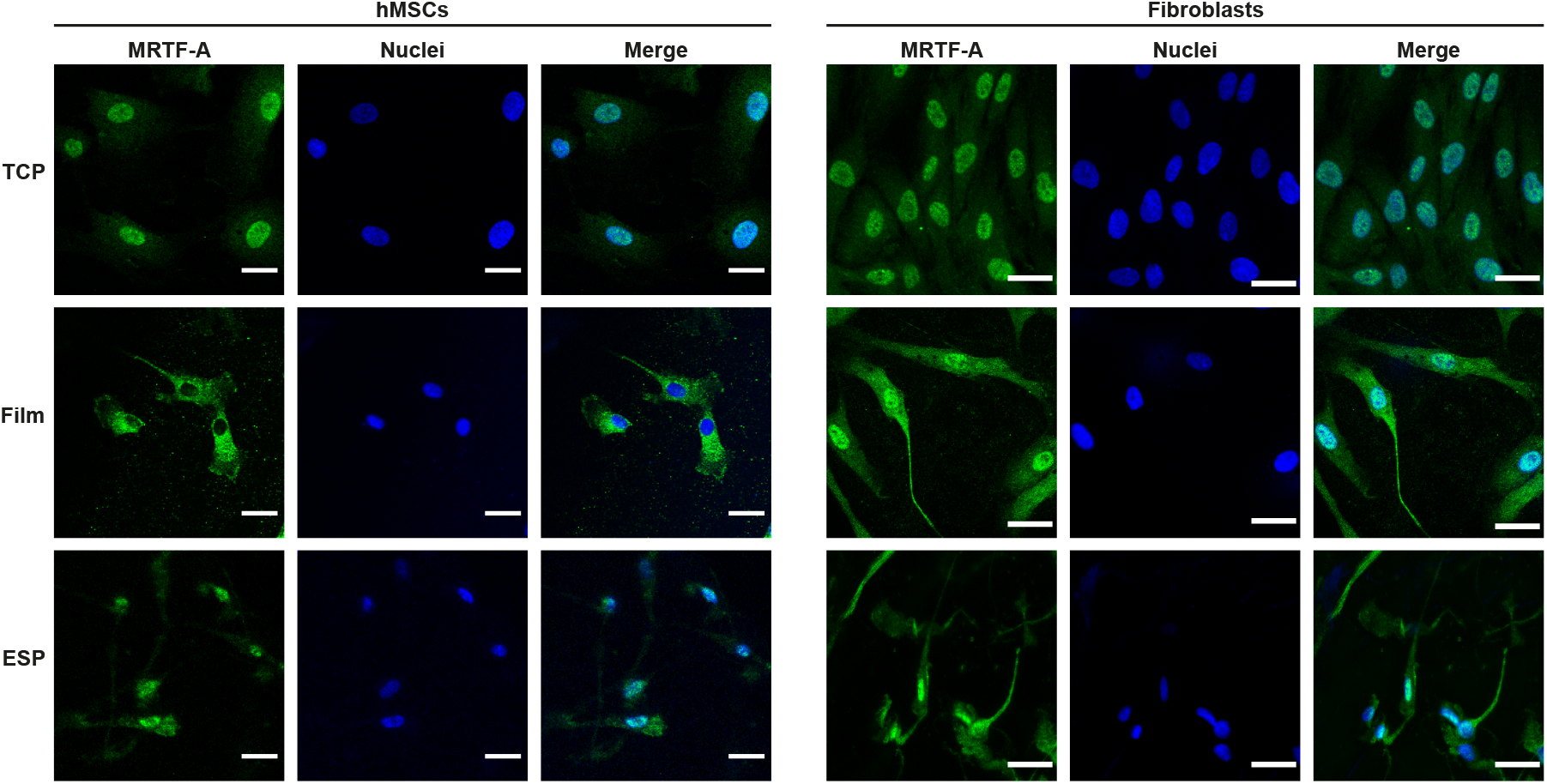
MRTF-A localization in hMSCs and fibroblasts on different culture substrates. hMSCs and fibroblasts were grown for 7 days on TCP, Films or ESP scaffolds and stained for MRTF-A (green) and nuclei (blue). Scalebars represent 30μm.

**Supplementary figure 6.**
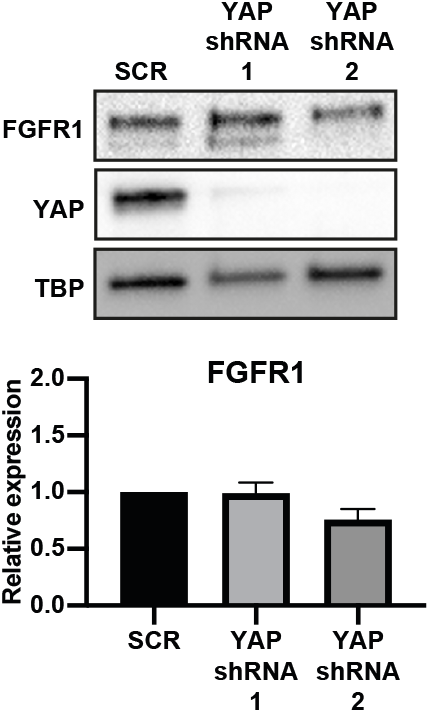
YAP does not regulate FGFR1 expression. **a, b,** Western blot of FGFR1, YAP and TBP (as loading control) of hMSCs transduced with YAP shRNA, cultured on TCP. Graphs depict quantification of western blots of FGFR1/TBP from 4 biological replica’s, normalized to TCP. Error bars indicate mean±SD. Repeated measures ANOVA with Tukey’s post-hoc test.

